# Biogeographic variation in reproductive strategy in a range-expanding marine gastropod

**DOI:** 10.1101/2025.09.24.678157

**Authors:** Keira S Monuki, Eric Sanford

## Abstract

Although species geographic distributions are increasingly shifting poleward with climate change, the processes that facilitate or impede range shifts are poorly understood. More extreme environmental conditions near poleward range boundaries often impose reproductive constraints, which may limit fitness and future range expansion. However, few studies evaluate the extent to which adaptive evolution and/or phenotypic plasticity might mitigate the effects of reproductive constraints and contribute to successful range shifts. We examined reproductive trait variation across the geographic range of *Acanthinucella spirata*, an intertidal snail that has recently expanded poleward along the coast of California, USA. We found that, compared to the range core, *A. spirata* from cooler range-edge populations allocated approximately 16x more nurse eggs to each developing embryo and had larger offspring at hatching, which may lead to greater juvenile growth rates and survival. Furthermore, range-edge *A. spirata* populations had lower fecundity, highlighting a potential tradeoff between offspring size and number. Some trait differences between range-core and range-edge populations persisted in an 18-month common-garden experiment, consistent with the hypothesis that such traits may be under selection. This study suggests that life-history adaptations in range-edge populations are an important and understudied mechanism that may contribute to species’ range expansions.

## Introduction

Climate change is leading to an unprecedented number of geographic range shifts as species distributions shift poleward to track warming temperatures [1–4]. Range shifts can greatly influence ecosystem dynamics, creating novel biotic communities and altering both genetic and species diversity [5–8]. As such, there is a strong interest in advancing our understanding of the processes that drive range expansions. Both theoretical and empirical work recognize the complexities of range shift dynamics and highlight a need to consider factors such as landscape context, demography, and evolutionary potential [9–11]. However, we still have incomplete knowledge of the ecological and evolutionary processes that facilitate or impede climate-driven range shifts. An understanding of the dynamics of range shifts is crucial to assess which species are likely to shift, and how quickly, with continued global change.

Studies of biogeography traditionally emphasize that extreme temperatures at the range edge interact with a species’ physiological tolerance limits to set its geographic distribution boundaries [12–14]. While suboptimal temperatures can limit a species’ range through lethal physiological effects, sublethal changes in temperature along latitudinal gradients can also have substantial impacts on performance and fitness [14]. In particular, reproductive constraints can occur when unfavorable temperatures restrict activity and energy gain, often leading to reduced stores of energy, slower growth, and reduced fecundity [14]. For example, cold temperatures can set poleward range limits by shortening summer growing seasons, failing to meet a temperature threshold to induce reproduction, and slowing or stopping juvenile development [12,13,15,16].

Range-edge environments that constrain reproduction, however, can also impose strong selection to counteract these effects through countergradient adaptations [17]. Selection imposed by cooler temperatures can lead to range-edge populations with increased rates of development, metabolism, growth, and fecundity relative to range core individuals, when measured at a common temperature [17,18]. Life-history tradeoffs may reinforce this countergradient trend towards increased growth and reproduction at the range edge. For example, if there are tradeoffs between investment in growth and reproduction versus competitive ability, low conspecific densities at the range edge are predicted to relax selection on competitive ability and favor traits that increase growth rates and reproduction [19,20]. Although past work suggests that biogeographic variation in reproductive constraints can be important for establishing range boundaries, we still know little about how such constraints might influence active range expansions.

Shifts in life-history strategies are one potential mechanism that may help counteract reproductive constraints during a range expansion as species move poleward into new environments. Studies of reproductive trait variation across latitude and elevation provide evidence that stressful environments can lead to life-history variation along an environmental gradient [21]. In particular, life histories can evolve through a trade-off between offspring number and offspring size (i.e., “many small vs. few large”) [22–24]. Studies show that females in higher elevation populations often produce fewer, higher quality offspring than those at lower elevations [25]. For example, dark-eyed juncos had fewer clutches at higher elevation but increased offspring survival, suggesting a life-history strategy shift towards increased parental investment in more stressful environments [26]. Plants show similar trends in offspring number vs. size in response to stress, where some species produce larger seeds under drought conditions [27] and exhibit a fecundity trade-off between seed number and size where fewer, larger seeds are favored in suboptimal environments [28]. There is also some evidence for life history shifts in species undergoing changes in geographic distribution. Hierro et al. (2020) found that seed size of the yellow star-thistle varied during an ongoing species invasion, where individuals had larger seeds in the invasive range compared to the range core, though other studies have not found a relationship between range expansion and seed size [29]. Therry et al. (2015) found that larger hatchling sizes in damselflies at a range expansion front resulted from variation in thermal regime across the range.

Variation in reproductive strategy during range expansion may arise from natural selection, phenotypic plasticity, or some combination of both [27,30]. Natural selection may play a substantial role in increasing reproductive success at the range edge [31,32]. In the northern tamarisk beetle, selection appears to have led to range-edge individuals with increased female body mass and fecundity [33]. Phenotypic plasticity may also help organisms acclimatize to more extreme environments at poleward range margins (34–36) and alter the spread of range expanding species [37,38]. While full plasticity would allow populations to respond to fine-scale environmental variation within one generation, this can be energetically costly, especially for small, resource-limited range edge populations in stressful and unpredictable environments [11]. Although there is some evidence for reproductive plasticity in response to extreme environments [27,34], few studies examine reproductive plasticity in species undergoing geographic distribution changes. Overall, the relative contributions of adaptation and plasticity remain an understudied component of range expansion dynamics.

In this study, we examined reproductive trait variation across the geographic range of the marine gastropod *Acanthinucella spirata*. *A. spirata* is a predatory intertidal whelk in the Family Muricidae that exhibits direct development with crawl-away juveniles [39]. The historic range of *A. spirata* spans from Bodega Bay, California, USA in the north to Baja California, Mexico in the south [4]. Recently, a sizable population of *A. spirata* was found in Cape Mendocino in northern California following marine heat events in 2014–2016 [40]. Although there have been some historical records of *A. spirata* in Mendocino County previously [4], no other populations have been seen north of Bodega Bay over the past few years [41]. In addition, observations suggest that *A. spirata* has experienced a striking increase in abundance in the Bodega Bay region during the last decade (E. Sanford, *personal communication*). Even with ongoing climate change, ocean temperature in the northern range edge of *A. spirata* is considerably cooler and more variable compared to the range core due to coastal upwelling (Figure S1).

Muricid gastropods, including *Acanthinucella* spp., reproduce by attaching egg capsules to rocky substrate [39]. Fertilized embryos typically develop within the capsules for 4-12 weeks, depending on the species, first passing through a larval veliger stage within the capsule and then hatching out as tiny juvenile snails that crawl away from the capsules [42]. Egg capsules of most muricid species also contain nurse eggs, which are non-developing eggs consumed by the developing embryos for nutrients [39]. The ratio of nurse eggs to developing embryos in muricid gastropods can vary among populations and among individual females [43,44]. Furthermore, greater investment in more nurse eggs per developing embryo leads to juvenile snails that hatch at larger sizes [43]. In turn, larger hatchling snails grow more quickly and have higher survival rates in the field [45]. Thus, we hypothesize that *A. spirata* may increase investment in nurse eggs as a strategy to increase their fitness in more extreme range-edge environments, especially as a species that lacks parental care behaviors [22].

Additionally, embryos and larvae of marine invertebrates develop more slowly under cooler temperatures, potentially increasing their exposure to predators and other sources of mortality [46,47]. To partially compensate for slower development rates under cooler temperatures, selection may favor countergradient variation in development times where embryos and larvae produced by higher-latitude females develop to hatching more quickly than those produced by conspecific females from lower latitudes, when held at a common temperature [48,49]. For example, Palmer (1994) found that development time in a high-latitude population of the muricid gastropod *Nucella ostrina* from Alaska was less sensitive to cooler temperatures than a population farther south in British Columbia.

The *A. spirata* range expansion provides the opportunity to test understudied questions regarding the importance of reproductive trait variation in facilitating or impeding ongoing geographic range shifts. Through this study, we investigate reproductive traits across populations of *A. spirata* to test the following hypotheses:

1. Reproductive traits vary across the geographic range of *A. spirata*. Relative to range-center populations, range-edge populations will have more nurse eggs per developing embryo, larger hatchling sizes, and decreased fecundity, consistent with the common tradeoff between offspring size and number.
2. When measured at a common temperature, individuals from range-edge populations will show faster development times and growth rates consistent with countergradient evolution.
3. *A. spirata* will rely primarily on phenotypic plasticity to adjust reproductive traits at the range edge to allow populations to respond quickly to highly variable range-edge environments.

## Methods

To investigate how reproductive traits vary across the range of a range-expanding species, we performed (1) a study of field-collected capsules to assess intraspecific reproductive trait variation across the geographic range of *Acanthinucella spirata*, and (2) a common garden laboratory experiment to assess the relative contributions of selection and plasticity to trait variation in egg capsules produced at two temperatures.

### Field-collected *A. spirata* egg capsules

#### Collections

We collected *A. spirata* egg capsules from five populations along the California coast from July to August 2023: Cabrillo Beach (CB), Rincon Point (RP), Muir Beach (MB), Dillon Beach (DB) and Cape Mendocino (CM) (Figure 1). We characterize Cabrillo Beach and Rincon Point as “range-core” populations and Muir Beach, Dillon Beach, and Cape Mendocino as “range-edge” populations where Cape Mendocino represents the current poleward extent of *A. spirata* (Table S1). At each site, we collected egg capsules from 7–10 distinct breeding aggregations. Within each aggregation, we collected 30–40 capsules that visually appeared early in development (i.e., bright yellow embryos and no visible shell formation). Capsules were brought back to the Bodega Marine Laboratory for analysis.

**Figure 1.**
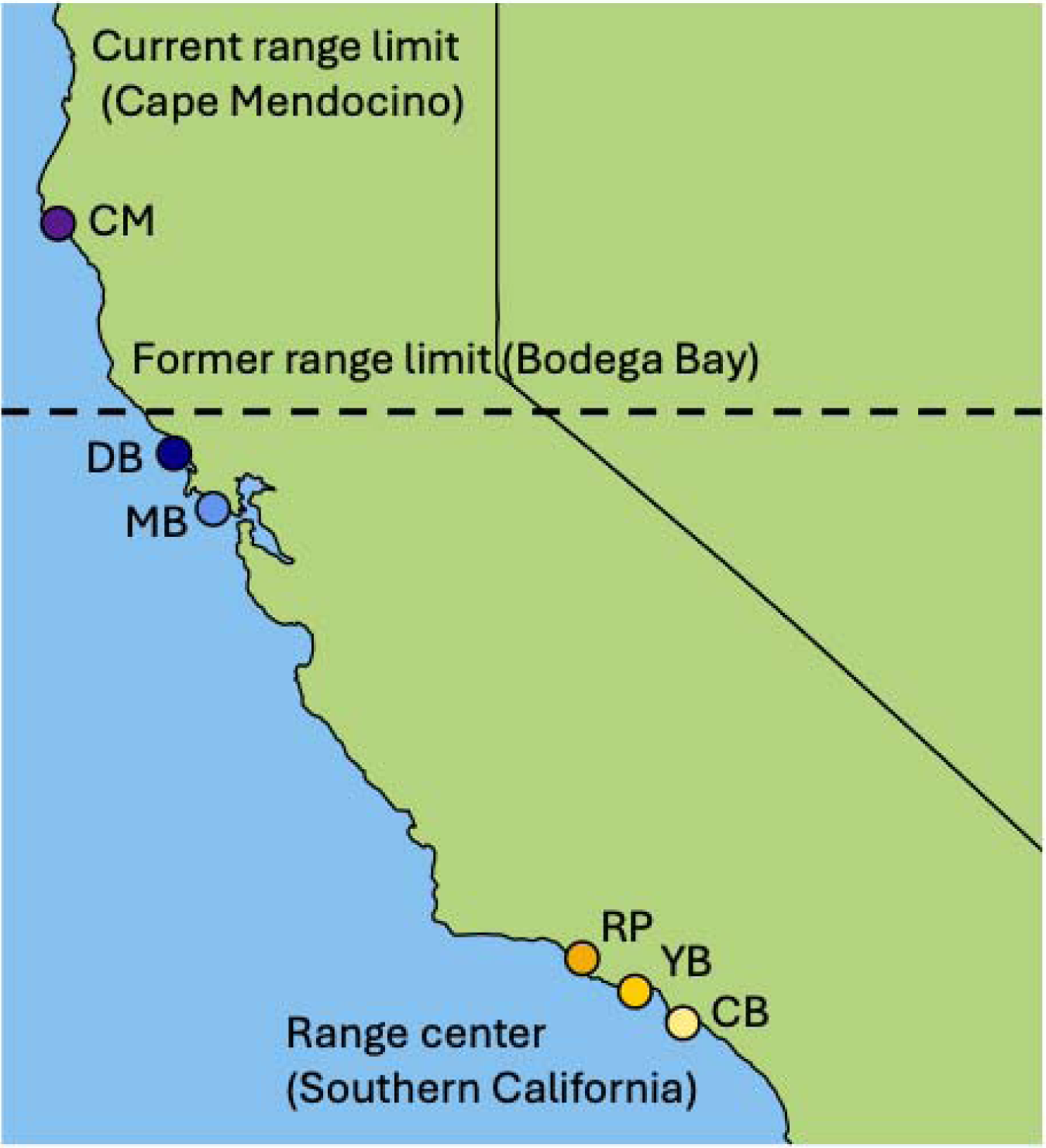
Map of study sites for the field collections and laboratory experiment for *Acanthinucella spirata* in California, USA. Darker colors (i.e., blues and purples) represent the range edge and lighter colors (i.e., yellows) represent the range core (Table S1). A dashed line marks the former range limit of Bodega Bay. Sites are abbreviated as follows: **CM** = Cape Mendocino (40°26’09.1“N 124°24’27.9“W); **DB** = Dillon Beach (38°15’17.0“N 122°58’12.6“W); **MB** = Muir Beach (37°51’28.0“N 122°34’27.3“W); **RP** = Rincon Point = (34°22’31.1“N 119°28’24.2“W); **YB** = Yerba Buena (34°03’09.8“N 118°57’51.2“W); **CB** = Cabrillo Beach (33°42’31.5“N 118°17’06.1“W).

#### Trait measurements

In the laboratory, we measured the following traits from the collected capsules for each aggregation: capsule length, number of embryos per capsule, embryo size, number of hatchlings per capsule, and hatchling size. We measured the length of the capsules by taking a photograph under a microscope (Leica M125 with camera attachment MC170 HD, Wetzlar, Germany) and measuring the length in ImageJ (version 1.53k, Schneider et al. 2012). We defined length as the horizontal distance between the bottom of the capsule, above the attachment site, to the point just proximal of the capsule plug (Figure S2A). We then haphazardly collected a subset of five capsules from each aggregation and dissected them to obtain embryo number and embryo size. Consumption of nurse eggs begins in the larval veliger stage and ends at hatching (Spight 1976a). As such, we only measured embryo traits for capsules in the pre-veliger development stage before nurse egg consumption began.

After dissecting the five capsules, we isolated three capsules from each aggregation and left them to develop until hatching in individual mesh-sided plastic tea strainers with ∼180 μm openings (Upton Tea Imports, Holliston, Massachusetts, USA). After hatching, we counted the number of hatchlings and measured the size of five haphazardly selected individuals from each capsule. We measured hatchling size by taking a photograph under a microscope and measuring the distance from the shell apex to the siphonal notch using ImageJ (Figure S2B). We estimated the ratio of nurse eggs to developing embryos using the following formula: 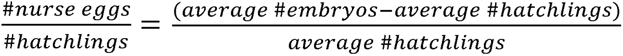 [39,51].

#### Statistical analyses

To analyze reproductive trait variation among populations, we used linear mixed effects and generalized linear mixed models (GLMM) with a log link function in the packages *lme4*, *stats* and *MASS* [52,53] in R (version 4.4.0; R Core Team, 2024). For capsule length, embryo number, and hatchling number, we used a GLMM with population as a fixed effect and aggregation as a random effect (Number ∼ Population + (1 | Aggregation)). For the hatchling size GLMM, we used population as a fixed effect and capsule as a random effect nested within the aggregation random effect (Size ∼ Population + (1 | Aggregation/Capsule)). For embryo size, we used a linear model with population as a fixed effect and capsule nested within aggregation as a random effect. We used a linear model with population as a fixed effect for nurse egg ratios. Model fit was assessed using residual histogram and Q-Q plots. We determined fixed effect significance using a Type II Wald chi-squared analysis of deviance using the Anova function in the *car* package [54]. All pairwise comparisons were conducted using the contrast function in *emmeans* [55].

### Common garden laboratory experiment

#### Collections and breeding experiment

We collected *A. spirata* individuals from four sites from December 2022 to January 2023: Yerba Buena (YB), Rincon Point (RP), Muir Beach (MB) and Dillon Beach (DB) (Figure 1). Yerba Buena and Rincon Point are range core sites, and Muir Beach and Dillon Beach are range edge sites (Table S1). We limited collections to sites where *A. spirata* were relatively common to minimize any negative impacts on small populations such as Cape Mendocino and Cabrillo Beach. At each site, we collected 150–200 snails whose lengths were between 12–17mm. *A. spirata* become reproductively mature at lengths greater than 17mm (K. Monuki, *personal observation*), so we aimed to collect immature snails to minimize the chances that prior exposure to field conditions would influence reproductive traits. We brought the snails back to the Bodega Marine Laboratory and divided them into two sump tanks at 11°C, which represents a typical low ocean temperature at the range edge and 16°C, which represents a typical average ocean temperature near the range core (Figure S1). Tanks were supplied with a continuous low flow of seawater and temperatures were maintained in the cool and warm treatments to within ± 1°C using a chiller (Aqualogic Cyclone, Model CY-3) and heater (500W titanium heater attached to a Model CN76120 controller, Omega Engineering, Inc.), respectively. Snails were fed local barnacle species (*Chthamalus dalli* and *Balanus glandula*) on small rocks collected from Campbell Cove in Bodega Harbor. We fed the snails *ad libitum* every two weeks over the course of 10 months until they reached reproductive maturity (>17mm) in late Fall 2023. Once the snails reached reproductive maturity, we determined their sex through visual inspection and placed them into breeding pairs consisting of one male and one female from the same population (Table S2). The breeding pairs (n = 34–39 per population) were then placed in separate 0.5L containers and subsequently moved from the tanks to a manifold setup. The manifold setup was a flow-through seawater system with temperature-controlled water in sump tanks flowing continuously through individual water lines with irrigation drippers to the 0.5L containers. We controlled sump tank temperatures using a 1000W titanium heater (Innovative Heat Concepts) and chiller (Aqualogic, Model DS-7) attached to controllers to maintain temperature at a set point of 12°C and 15°C, respectively (Figure S3). Water temperature was generally maintained within ± 1°C. The replicate containers were placed in a random order across the manifold setup, water flowed through replicate containers only once, and no water was shared among the containers to maintain their independence. We placed breeding pairs into the manifold setup in November 2023, or in January 2024 for individuals that grew more slowly (Table S2). Containers were cleaned regularly (1–2x a week), and snails were supplied with fresh rocks with barnacles every two weeks until the beginning of the reproductive season in March 2024. During the reproductive season, we fed the snails once a month to minimize disturbance during egg laying.

#### Trait measurements

For the common garden experiment, we measured the same traits from the study of field-collected capsules (capsule length, embryo number, embryo size, hatchling number, hatchling size), using the same methods described above (see section “Trait measurements”). We also measured hatchling development time, nurse egg ratios per breeding female, fecundity of the breeding females, and absolute growth rate 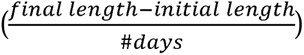 of all adult snails. To quantify hatchling development time, we recorded the date of the first capsule laid by the breeding pair. We then isolated three capsules in individual tea strainers and let them develop through hatching. We measured the end of development as the date when the first juvenile snail escaped from the capsule plug. We then estimated nurse egg ratios per breeding female using the same equation described above for the field-collected capsules [39,51]. We estimated fecundity as the number of laid capsules multiplied by the average number of hatchlings per capsule for each breeding female. We also measured the shell lengths of the adult snails, defined as the length between the shell apex to the siphonal notch, using handheld calipers at the beginning and end of the experiment to quantify differences in growth across temperature treatments and across populations (Figure S4).

#### Statistical analyses

To analyze trait variation across populations, we used linear mixed effects models and GLMMs with a log link function, population and female size as fixed effects, and breeding pair as a random effect (Methods S1). Because only three snails laid eggs in the 12°C treatment, we excluded the cold treatment from the reproductive trait analyses (Methods S2). Similarly, only two females from Rincon Point laid eggs, so we excluded the Rincon Point population from the reproductive trait analyses. We included female size as a covariate in the models because, although females that produced capsules were within a relatively small size range (shell lengths = 18.5–25.3 mm), there was a positive relationship between female size and some reproductive traits (Figure S5). Model fit was assessed using residual histogram and Q-Q plots. Significance and pairwise comparisons were conducted using the same methods as the field study.

## Results

Overall, reproductive traits varied strongly across populations. In general, range-edge populations had larger capsules, more embryos per capsule, greater nurse egg ratios and larger hatchlings in the field. In contrast, range-core populations had higher fecundity. Some, but not all, traits differences persisted in common garden conditions.

### Field-collected capsules

Capsule length varied among populations (Table S3; GLMM, gamma, “population”, χ2 = 112.53, p<0.001). The two northernmost sites Dillon Beach and Cape Mendocino had the largest capsules and range-core site Cabrillo Beach had the smallest capsules (Figure 2A). Range-core site Rincon Point and range-edge site Muir Beach had intermediate sized capsules (Figure 2A). Embryo number also varied among populations (Table S4; GLMM, neg binomial, “population”, χ2 = 54.63, p<0.001). Range-edge sites Muir Beach, Dillon Beach, and Cape Mendocino egg capsules had approximately 48% more embryos per capsule than those from range-core sites Cabrillo Beach and Rincon Point (Figure 2B). Cabrillo Beach had larger embryos than Rincon Point, Muir Beach, and Dillon Beach, but not Cape Mendocino (Figure 2C; Table S5; linear mixed model, “population”, χ2 = 15.727, p = 0.003). Hatchling number did not vary among populations (Figure 2D; GLMM, neg binomial, “population”, χ2 = 5.8352, p = 0.2118). Capsules from range-edge sites Muir Beach and Dillon Beach had approximately 16x more nurse eggs per developing embryo than capsules from range-core site Cabrillo Beach, and range-core site Rincon Point and range-edge site Cape Mendocino capsules had intermediate nurse egg ratios (Figure 2E; Table S6; linear model, “population”, F_4,17_ = 8.2905, p<0.001). Hatchling size varied among populations (Table S7; GLMM, gamma, “population”, χ2 = 75.74, p<0.001). The two northernmost sites Dillon Beach and Cape Mendocino had larger hatchlings than Cabrillo Beach, Rincon Point and Muir Beach (Figure 2F).

**Figure 2.**
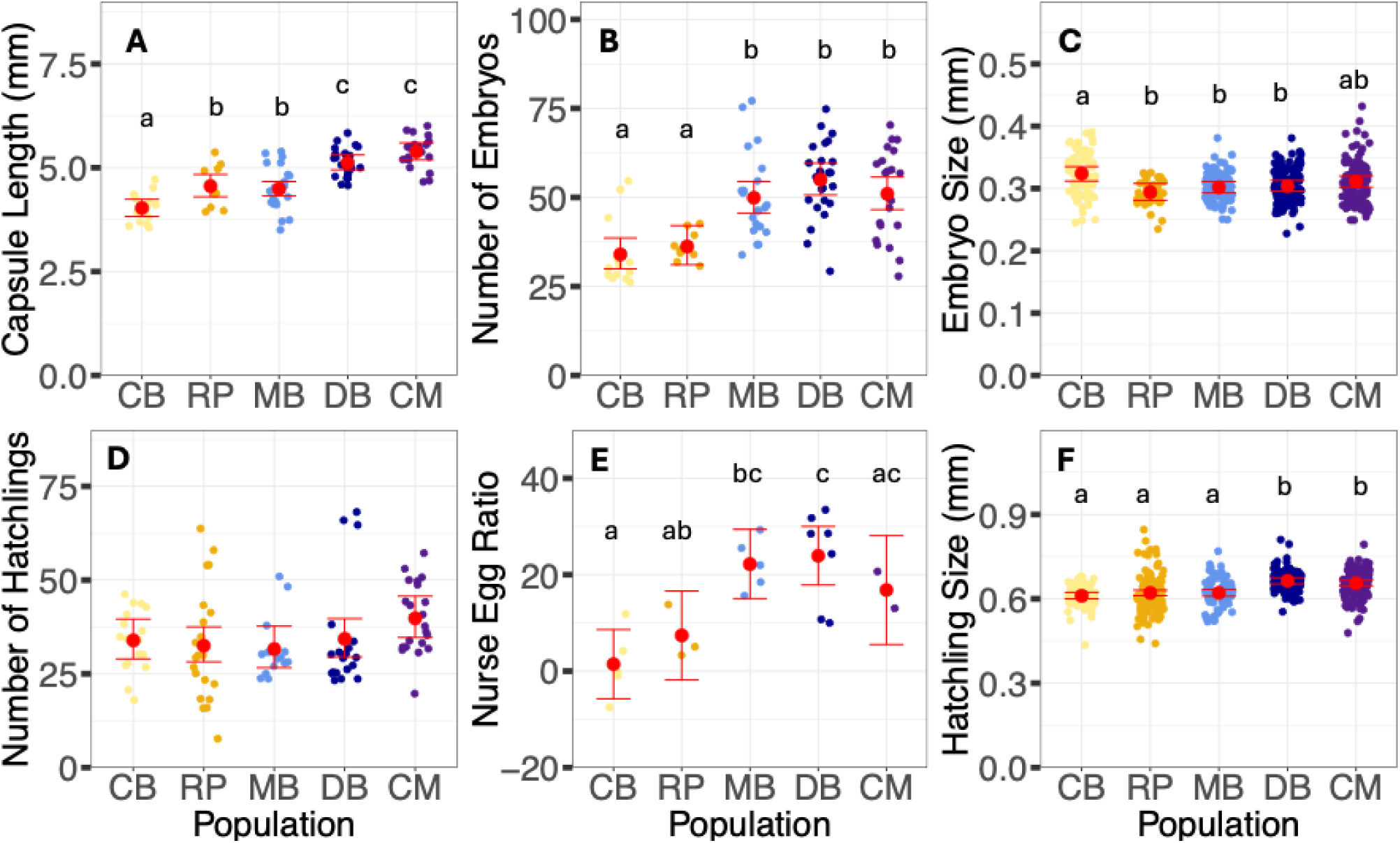
Variation in reproductive traits across the geographic range of *Acanthinucella spirata* for field-collected capsules. **(A)** egg capsule length, **(B)** number of embryos per capsule, **(C)** embryo size, **(D)** number of hatchlings per capsule, **(E)** nurse egg ratio estimates per breeding aggregation and **(F)** hatchling size. Populations are ordered South to North on the x-axis and use the same site abbreviations and color scheme as Figure 1. The raw data are scattered for each population. The population-level model estimates and 95% confidence intervals are back-transformed from the log scale and are shown in red. Different letters represent significant differences between pairwise comparisons (p<0.05).

### Common garden experiment

We obtained egg capsules from 34 breeding pairs in the 15°C treatment (n=12 from Yerba Buena, n=11 from Muir Beach, n=11 from Dillon Beach). We obtained hatchling measurements from 31 breeding pairs, as capsules from three breeding pairs did not hatch. Only two breeding pairs from Rincon Point laid egg capsules at 15°C, and only three breeding pairs in the 12°C treatment laid capsules. As such, we only report results for the three populations (Yerba Buena, Muir Beach, and Dillon Beach) in the 15°C treatment.

Some, but not all, of the results from the lab experiment generally mirrored those from analyses of field-collected capsules. Range-edge sites Dillon Beach and Muir Beach had smaller embryos than range-core site Yerba Buena (Figure 3C; Table S8; linear model, “population”, χ2 = 7.0694, p = 0.02917). Hatchling size varied among populations (Table S9; GLMM, gamma, “population”, χ2 = 20.7863, p<0.001), and Dillon Beach had significantly larger hatchlings than both Muir Beach and Yerba Buena (Figure 3F). Hatchling number did not vary across populations (Figure 3D; GLMM, neg binomial, “population”, χ2 = 1.8955, p = 0.3876). Although the number of embryos per capsule (GLMM, neg binomial, “population”, χ2 = 2.7399, p = 0.2541) and nurse egg ratios (linear mixed model, “population”, F_2,29_ = 1.4871; p = 0.2428) did not vary significantly among populations, they showed similar qualitative trends where range-edge sites Dillon Beach and Muir Beach tended to have more embryos per capsule and higher nurse egg ratios than range-core site Yerba Buena (Figure 3B and 3E). Contrary to the field-collected capsules, range-edge site Dillon Beach had smaller capsules than range-core site Yerba Buena (Figure 3A; Table S10; GLMM, gamma, “population”, χ2 = 6.053, p = 0.04848). Development time (Figure 4A; GLMM, neg binomial, “population”, χ2 = 3.0217, p = 0.22172) did not differ among populations. Fecundity varied across populations (GLMM, gamma, “population”, χ2 = 7.4695, p = 0.02388), where range-edge site Dillon Beach had significantly lower fecundity than range-core site Yerba Buena, and range-edge site Muir Beach had intermediate fecundity (Figure 4B; Table S11).

**Figure 3.**
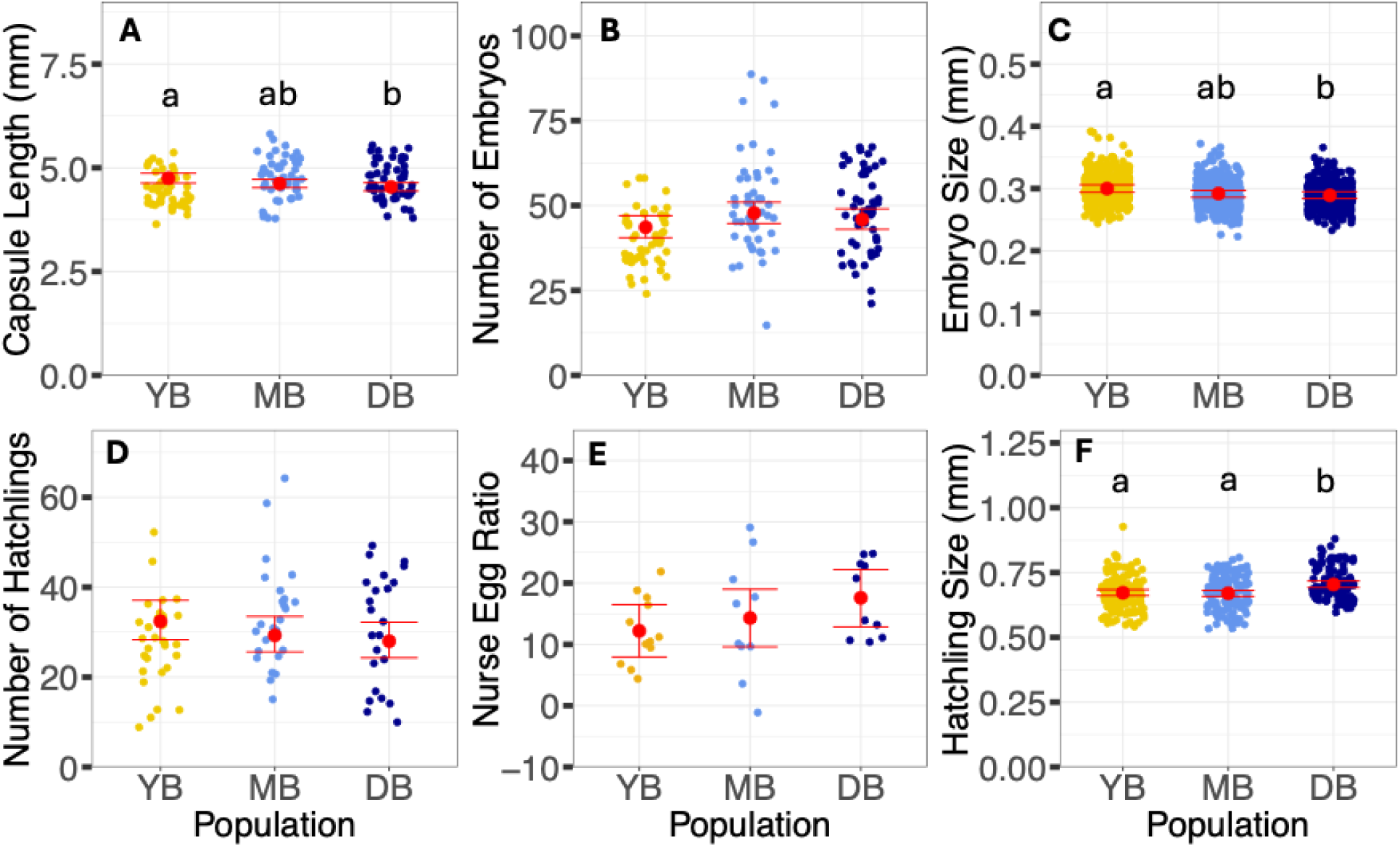
Variation in reproductive traits among populations of *Acanthinucella spirata* for capsules produced in the common garden experiment. **(A)** egg capsule length, **(B)** number of embryos per capsule, **(C)** embryo size, **(D)** number of hatchlings per capsule, **(E)** nurse egg ratio estimates per breeding aggregation and **(F)** hatchling size. Populations are ordered South to North on the x-axis and use the same abbreviations and color scheme as Figure 1. The raw data are scattered for each population. The population-level model estimates and 95% confidence intervals are back-transformed from the log scale and are shown in red. Different letters represent significant differences between pairwise comparisons (p<0.05).

**Figure 4.**
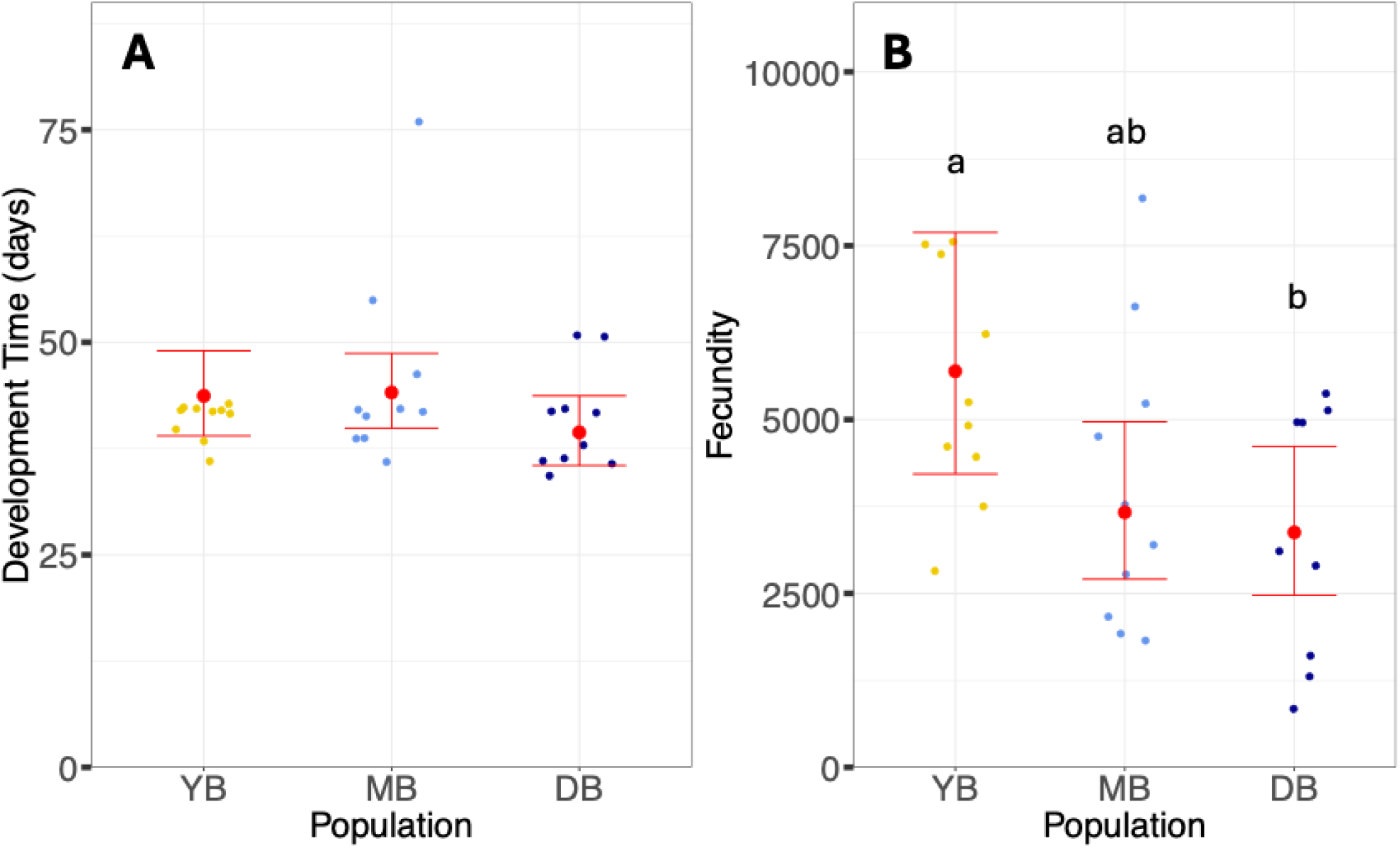
Variation in reproduction among populations of *Acanthinucella spirata* in the common garden experiment. **(A)** development time in days **(B)** fecundity (estimated total number of hatchlings per female = number of capsules*average number of hatchlings per capsule). Populations are ordered South to North on the x-axis and use the same abbreviations and color scheme as Figure 1. The raw data are scattered for each population. The population-level model estimates and 95% confidence intervals are back-transformed from the log scale and are shown in red. Different letters represent significant differences between pairwise comparisons (p<0.05).

Adult snail growth rates were not significantly different across populations (linear model, “population”, F_3,258_= 2.3289, p = 0.075). Growth rate did vary by temperature treatment (linear model, “treatment”, F_1,258_ = 7.6774, p = 0.006) and by sex (linear model, “sex”, F_1,258_ = 24.7351, p<0.001). Snails in the 12°C treatment overall grew faster than snails in the 15°C treatment and female snails grew faster than male snails (Figure 5). For range-core site Rincon Point, however, snails in the 15°C treatment grew faster than snails in the 12°C treatment (Figure 5; linear mixed model, “population*treatment”, F_3,258_ = 6.1578, p<0.001).

**Figure 5.**
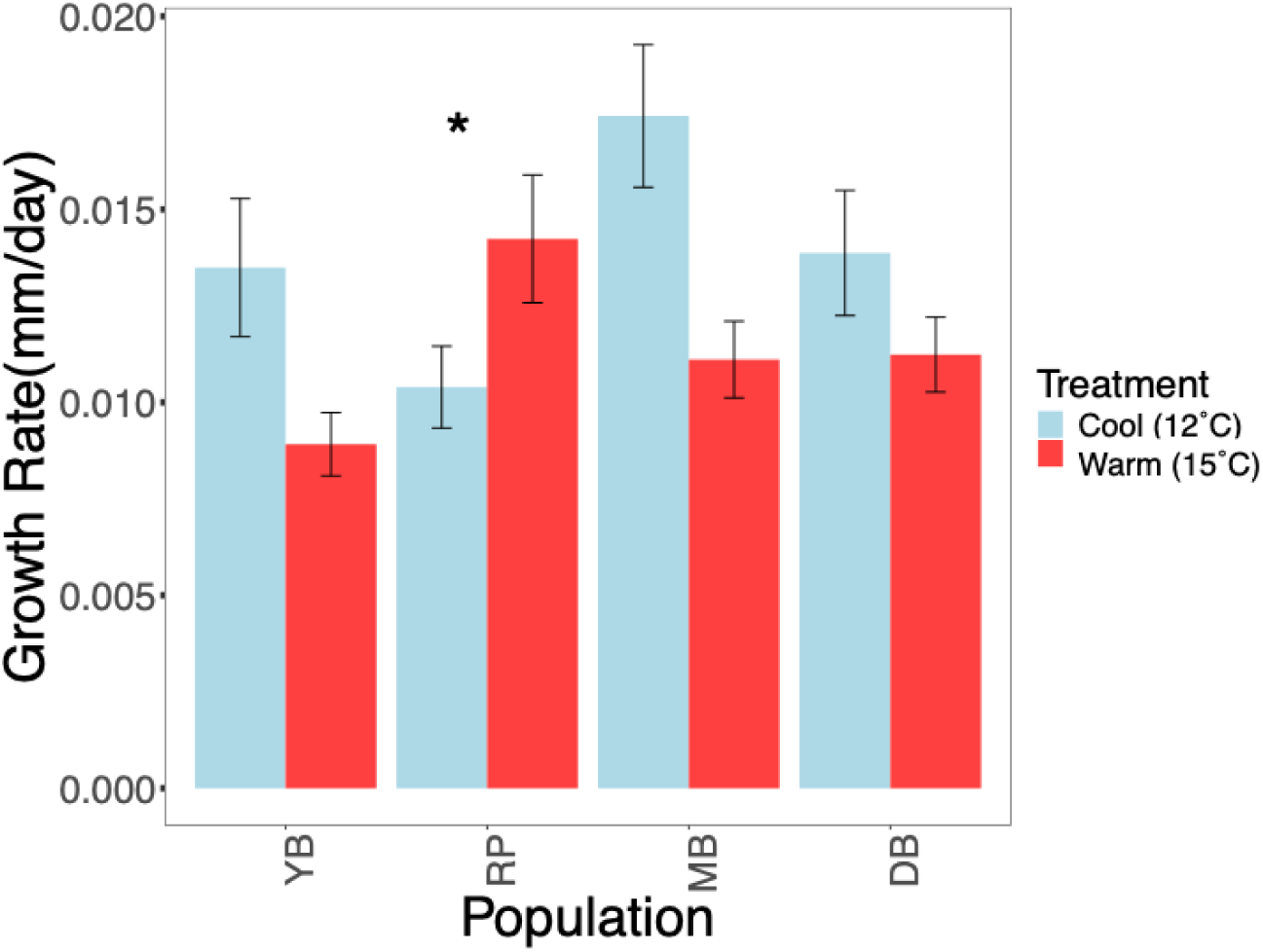
Growth rate of adult breeding snails (*Acanthinucella spirata*) from the common garden experiment. Populations are ordered from South to North on the x-axis and absolute growth rate (mm/day) is on the y-axis. Mean and standard error bars are shown for each population and for each temperature treatment. The cool treatment is 12°C and the warm treatment is 15°C. Asterisk indicates the significant Population*Treatment interaction (F_3,258_ = 6.1578, p<0.001), as Rincon Point (RP) was the only population with greater growth in the warm treatment than the cool treatment.

## Discussion

Through this study, we investigated reproductive trait variation in the ongoing range shift of *Acanthinucella spirata* and explored potential contributions of selection and plasticity to such variation. Range-edge populations of *A. spirata* had a suite of reproductive traits that were strikingly different from the range core. Specifically, range-edge populations had more embryos per capsule, greater ratios of nurse eggs per developing embryo, and larger hatchling sizes in the field. These traits are expected to confer higher population fitness in the face of colder temperatures near the range edge and might provide a mechanism by which *A. spirata* can respond to marginal poleward conditions during range expansion. Greater nurse egg ratios in muricid snails provide increased nutrition to developing larvae within egg capsules, leading to larger hatchling sizes and faster juvenile growth rates [44,45]. In addition to having faster growth as juveniles, larger hatchling snails have higher survival [45]. Evidence for the fitness advantage of larger hatchlings are seen in other aquatic taxa such as crustaceans and turtles, as larger hatchlings have higher tolerances for unfavorable conditions, can escape size-specific predation, and can outcompete smaller conspecifics [56,57].

Given the association between increased nurse egg ratios and larger hatchling sizes, we hypothesize that this shift toward greater nurse egg allocation at the range edge may increase juvenile performance and contribute to the continued poleward expansion of *A. spirata*. In particular, females that allocate greater nutritional investment to each offspring may increase reproductive success. This life-history strategy is a common adaptation documented in other taxa, such as birds and plants, to increase fitness in more extreme environments near species’ distributional limits [25,58]. Furthermore, fecundity was lower and hatchling sizes were larger at the range edge, highlighting a potential offspring size vs. number tradeoff where range-edge females invest more energy into fewer, larger offspring [22,59]. Despite this tradeoff, selection that favors shifts in reproductive strategy in novel environments may help facilitate poleward range expansions if it ultimately leads to greater survival of range-edge offspring.

We found evidence that some reproductive trait differences in range-edge populations may be a result of selection. Embryo and hatchling size variation was similar in both the field-collected and laboratory-produced capsules, even though the common garden study was conducted in a controlled environment where snails experienced the same conditions for ∼18 months. Some traits that did not vary significantly in laboratory-produced capsules (i.e., number of embryos, nurse egg ratios) showed qualitatively similar trends to the field-collected capsules (Figure 3B, E). As such, our results are broadly consistent with the hypothesis that greater nurse egg allocation and larger hatchling sizes are adaptations resulting from selective pressures at the range edge. These results, however, do not completely eliminate plasticity as a potential driver of range-wide variation as the patterns in reproductive trait variation in the field did not exactly match the patterns in the common garden experiment. Additionally, while we tried to minimize exposure to field conditions in our common garden experiment by collecting *A. spirata* when they were small and reproductively immature, there still may have been unknown environmental cues in early life or maternal inheritance (i.e., transgenerational plasticity) that influenced reproductive trait values [60]. Studies of range-shifting species with shorter generation times may help tease apart the relative importance of adaptation and plasticity in facilitating range shifts [61].

Environmental variation at the range edge may lead to shifts in the importance of adaptation versus phenotypic plasticity [36]. In general, plasticity is hypothesized to be important in highly variable environments, whereas stable environments tend to promote selection [62]. Therefore, one might expect that highly variable range-edge environments favor plasticity during range expansion [63]. However, our finding that range edge populations produce larger hatchlings even in common-garden conditions suggests that selection is, at least partially, shaping trait variation during the *A. spirata* range shift. One explanation is that local adaptation is often important for species with benthic development and low dispersal potential, like *A. spirata* [64]. Another explanation is that the short timescale of variability (i.e., days to weeks) in range-edge environments associated with episodic upwelling events in Northern California leads to selection to minimize nonadaptive plasticity. Past studies suggest nonadaptive plasticity may be more common than adaptive plasticity in novel environments [62,65,66]. Therefore, if the environment varies on a timescale where parental environment during egg capsule production differs from the offspring environment, when snails hatch 7–8 weeks later, directional selection that reduces nonadaptive plasticity may be favored [62,66]. Selection may also be important if there are high energic costs associated with plasticity in environments with high variability [11]. Our findings are consistent with the hypothesis that selection drives reproductive trait variation if range-edge environments are cooler and more variable on relatively short timescales (Figure S1).

We found variation in adult growth rates between the two temperature treatments, although not between populations. In general, snails grew faster in the cold treatment than the warm treatment. These results are counter to our expectations based on countergradient selection [17] and general metabolic theory, in which higher temperatures should lead to faster metabolism and greater growth, depending on where the temperatures experienced lie along the thermal performance curve [67]. It is possible that *A. spirata* alter their resource allocation strategy depending on the thermal environment. For example, snails may prioritize somatic growth over reproduction in cold environments, or perhaps conditions that are too cold to support successful reproduction may mean snails have more energy to devote to growth. Conversely, in less stressful warm conditions, snails may allocate more energy towards reproduction at the expense of slower growth. In this study, snail populations in the cold treatment did not lay eggs and generally had elevated growth rates, supporting the hypothesis that adult snails may allocate energy towards growth in cooler temperatures. We also saw evidence for genotype by environment (GxE) interactions in growth rate, where the relationship between growth rate and temperature varied depending on population of origin [68]. In contrast to other populations, *A. spirata* from range core site Rincon Point grew faster in the warm treatment than the cold treatment, consistent with expectations of local adaptation. This population-level variation indicates there may be some genetic basis to growth rate that selection can act upon, which may be important for future range expansion.

Our study provides an important empirical example of increased investment in individual offspring at the range edge and an offspring size vs. number tradeoff during a range expansion. With accelerating global change, consideration of how selection and plasticity may operate on reproductive traits across the range of a shifting species is critical to understand the processes that facilitate or impede climate-driven range shifts.

