## Supplemental Materials for "Biogeographic variation in reproductive strategy in a range-expanding marine gastropod"

**Methods S1.**

For capsule length, embryo number, and hatchling number, we used a GLMM with female size and population as fixed effects and breeding pair as a random effect (Number ~ Female Size + Population + (1 | Breeding Pair)). For the hatchling size GLMM, we used female size and population as fixed effects and capsule as a random effect nested within the breeding pair random effect (Size ~ Female Size + Population + (1 | Breeding Pair/Capsule)). For embryo size, we used a linear model with female size and population as fixed effects and capsule nested within breeding pair as a random effect. We used a linear model with female size and population as fixed effects for nurse egg ratios. For development time and fecundity, we used a GLMM with female size and population as fixed effects. For absolute growth rate, since there was no relationship between initial size and absolute growth rate (Figure S6), we used a linear model with an interaction term for population * treatment * sex (Absolute Growth Rate ~ Population * Treatment * Sex). We included all individuals in the experiment, including those from the 12˚C treatment and Rincon Point population, in the growth analyses.

**Methods S2.**

We made various unsuccessful attempts to induce reproduction in the 12˚C treatment, in case a warm temperature pulse was necessary to begin egg laying. The first was a 6-hour ambient air exposure to simulate a late morning low tide on August 12, 2024. The second was a 24-hour warm-water exposure on August 19, 2024, where we turned off the chiller and allowed the water to reach ambient temperature for 24 hours. We did not want the snails to experience warm water for more than 24 hours in order to maintain the temperature treatments for the experiment.

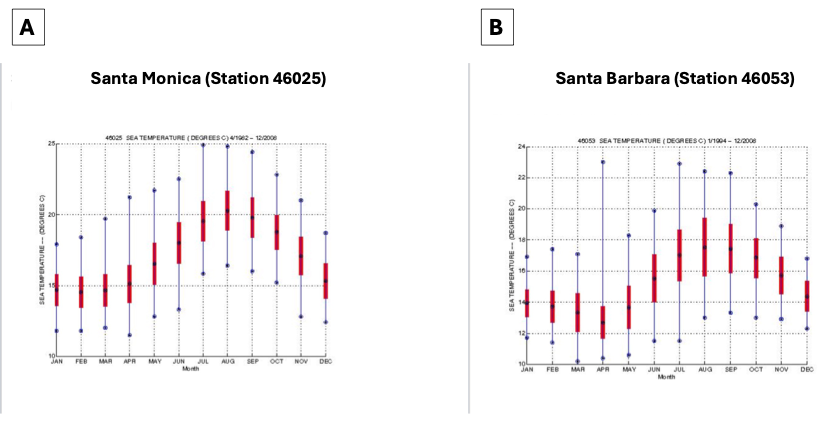

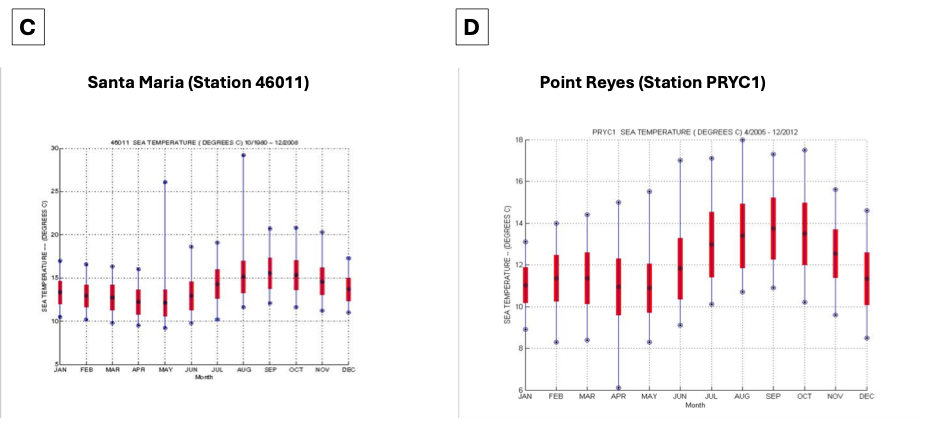

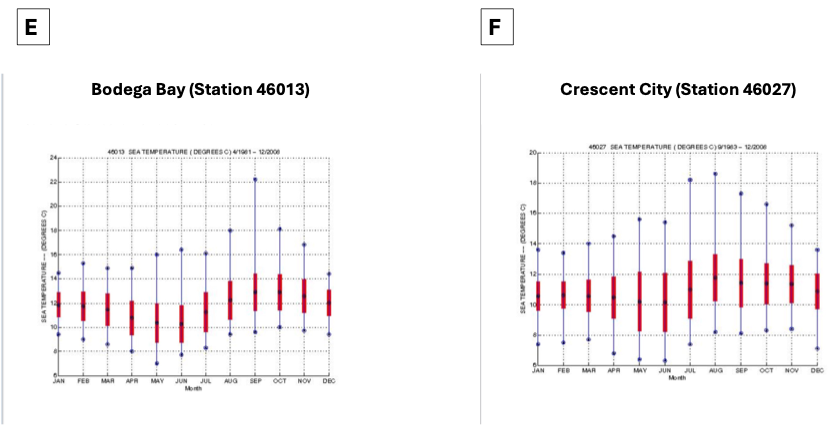

**Figure S1.** Sea surface temperature buoy data from the National Data Buoy Center and NOAA National Ocean Service Water Level Observation Network, showing means and standard deviations for each month based on data collected from 1980-2012. **(A)** Santa Monica (33°45'19" N 119°2'42" W; 50 km N of Cabrillo Beach) **(B)** Santa Barbara (34°14'26" N 119°50'20" W; 22 km N of Rincon Point) **(C)** Santa Maria (34°56'14" N 120°59'58" W) **(D)** Point Reyes (37°59'46" N 122°58'36" W; 35 km N of Muir Beach) **(E)** Bodega Bay (38°14'5" N 123°19'1" W; 26 km N of Dillon Beach) **(F)** Crescent City (41°50'24" N 124°22'54" W; 196 km N of Cape Mendocino)

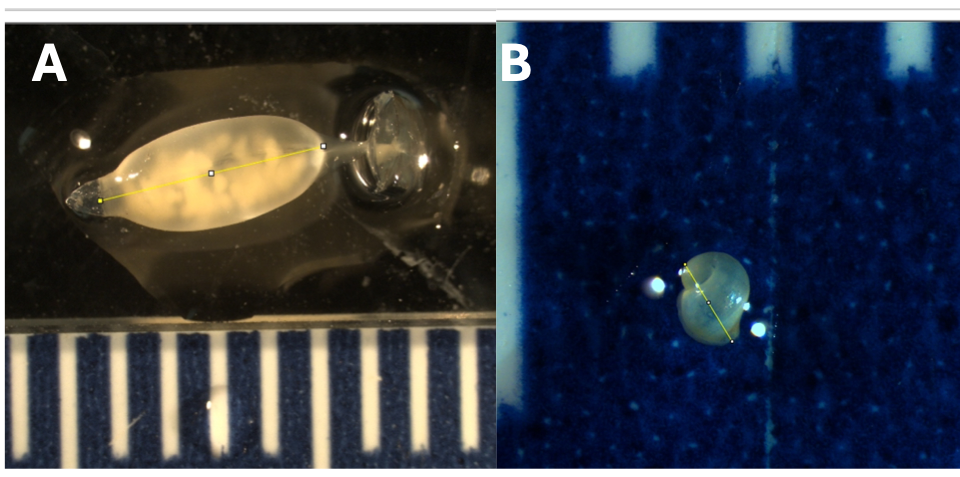

**Figure S2.** Measurements of reproductive traits in *Acanthinucella spirata*. **(A)** Image of egg capsule length measurement, and **(B)** hatchling length measurement in ImageJ. Scale bars are in mm.

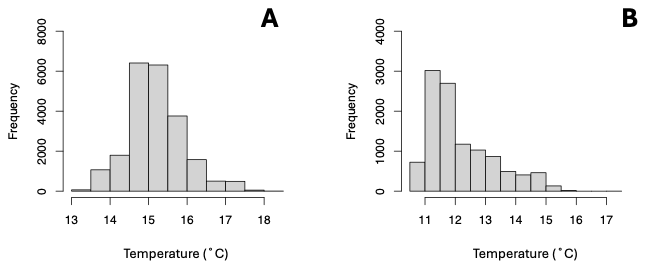

**Figure S3.** Histogram of temperatures (˚C) for two 0.5L containers in the **(A)** heated (mean = 15.18˚C) and **(B)** chilled (mean = 12.17˚C) manifold setup. TidbiT dataloggers recorded temperature every 30 minutes from 01/15/24 – 08/31/24.

| Length (mm) | Buoyant Weight (g) |
| --- | --- |
| 17.68 | 0.47 |
| 23.85 | 1.02 |
| 14.05 | 0.28 |
| 19.3 | 0.49 |
| 21.21 | 0.75 |
| 13.29 | 0.23 |
| 22.55 | 0.62 |
| 16.94 | 0.36 |
| 19.95 | 0.51 |
| 14.79 | 0.28 |
| 19.24 | 0.55 |
| 18.34 | 0.51 |
| 22.75 | 0.85 |
| 20.49 | 0.62 |
| 19.31 | 0.56 |
| 11.15 | 0.15 |
| 16.08 | 0.35 |
| 13.71 | 0.23 |
| 15.88 | 0.3 |
| 17.25 | 0.46 |
| 16.41 | 0.38 |
| 17.07 | 0.46 |
| 17.33 | 0.46 |
| 16.24 | 0.42 |
| 18.06 | 0.48 |
| 17.31 | 0.48 |
| 14.05 | 0.26 |
| 17.46 | 0.46 |
| 17.62 | 0.5 |
| 15.61 | 0.28 |
| 16.12 | 0.36 |
| 18.62 | 0.56 |
| 18.38 | 0.46 |
| 16.84 | 0.36 |
| 18.38 | 0.45 |
| 19.19 | 0.59 |
| 13.91 | 0.26 |
| 20.63 | 0.81 |
| 15.9 | 0.5 |
| 16.74 | 0.38 |
| 15.18 | 0.35 |
| 18.15 | 0.51 |
| 14.8 | 0.38 |
| 19.67 | 0.67 |
| 14.87 | 0.32 |
| 17.6 | 0.48 |
| 17.23 | 0.47 |
| 16.23 | 0.34 |
| 17.33 | 0.54 |
| 13.77 | 0.24 |
| 15.23 | 0.4 |
| 14.1 | 0.26 |
| 16.96 | 0.56 |
| 17.02 | 0.49 |
| 16.21 | 0.5 |
| 15.36 | 0.41 |
| 16.69 | 0.52 |
| 14.52 | 0.31 |
| 14.76 | 0.33 |
| 18.39 | 0.52 |
| 19.38 | 0.61 |
| 16.82 | 0.42 |
| 16.55 | 0.41 |
| 14.41 | 0.3 |
| 21.3 | 0.86 |
| 16.32 | 0.42 |
| 14.16 | 0.31 |
| 14.8 | 0.29 |
| 15.15 | 0.34 |
| 16.97 | 0.44 |
| 15.92 | 0.34 |
| 16.46 | 0.28 |
| 13.87 | 0.26 |
| 16 | 0.3 |
| 13.45 | 0.23 |
| 14.84 | 0.33 |
| 13.56 | 0.19 |
| 16.85 | 0.52 |
| 16.03 | 0.39 |
| 12.03 | 0.18 |
| 17.85 | 0.52 |
| 22.61 | 0.71 |
| 21.16 | 0.82 |
| 21.87 | 0.83 |
| 19.39 | 0.53 |
| 17.33 | 0.46 |
| 18.42 | 0.55 |
| 18.99 | 0.52 |
| 19.75 | 0.61 |
| 16.88 | 0.48 |
| 19.39 | 0.63 |
| 18.52 | 0.51 |
| 23.55 | 1.06 |
| 17.56 | 0.56 |
| 22.44 | 0.91 |
| 16.58 | 0.43 |
| 16.84 | 0.48 |
| 17.29 | 0.48 |
| 18.13 | 0.53 |
| 15.4 | 0.39 |
| 16.73 | 0.41 |
| 14.54 | 0.32 |
| 18.64 | 0.66 |
| 16.23 | 0.43 |
| 13.07 | 0.25 |
| 15.49 | 0.35 |
| 16.68 | 0.41 |
| 14.2 | 0.29 |
| 14.25 | 0.31 |
| 14.33 | 0.31 |
| 14.84 | 0.3 |
| 12.97 | 0.22 |
| 14.4 | 0.31 |
| 17.51 | 0.47 |
| 15.01 | 0.31 |
| 15.49 | 0.29 |
| 17.37 | 0.38 |
| 17.28 | 0.38 |
| 15.39 | 0.31 |
| 16.28 | 0.4 |
| 12.87 | 0.22 |
| 19.2 | 0.53 |
| 17.33 | 0.42 |
| 16.51 | 0.37 |
| 16.71 | 0.55 |
| 13.89 | 0.28 |
| 18.44 | 0.51 |
| 18.6 | 0.45 |
| 15.79 | 0.31 |
| 15.73 | 0.41 |
| 16.22 | 0.39 |
| 9.74 | 0.09 |
| 17.38 | 0.48 |
| 13.7 | 0.25 |
| 12.34 | 0.19 |
| 11.88 | 0.17 |
| 15.07 | 0.34 |
| 14.51 | 0.29 |
| 17.98 | 0.42 |
| 14.71 | 0.33 |
| 11.01 | 0.17 |
| 15.21 | 0.26 |
| 16.27 | 0.31 |
| 13.84 | 0.22 |
| 18.87 | 0.58 |
| 20.03 | 0.55 |
| 18.5 | 0.57 |
| 18.74 | 0.59 |
| 19.9 | 0.52 |
| 13.76 | 0.23 |
| 15.09 | 0.3 |
| 15.14 | 0.25 |
| 14.6 | 0.24 |
| 22.55 | 0.65 |
| 18.35 | 0.55 |
| 17.87 | 0.48 |
| 16.13 | 0.31 |
| 15.76 | 0.39 |
| 17.61 | 0.51 |
| 15.59 | 0.34 |
| 14.03 | 0.24 |
| 17.45 | 0.67 |
| 14.15 | 0.26 |
| 13.41 | 0.27 |
| 19.76 | 0.68 |
| 16.87 | 0.43 |
| 18.22 | 0.72 |
| 17.99 | 0.53 |
| 17.07 | 0.36 |
| 13.68 | 0.23 |
| 17.15 | 0.53 |
| 19.5 | 0.72 |
| 15.62 | 0.39 |
| 22.21 | 0.75 |
| 21.19 | 0.72 |
| 11.74 | 0.18 |
| 16.22 | 0.46 |
| 17.49 | 0.54 |
| 13.53 | 0.28 |
| 17.7 | 0.51 |
| 19.19 | 0.61 |
| 19.57 | 0.7 |
| 17.66 | 0.51 |
| 18 | 0.53 |
| 21.32 | 0.94 |
| 14.29 | 0.35 |
| 16.38 | 0.43 |
| 18.38 | 0.68 |
| 19.36 | 0.59 |
| 19.46 | 0.65 |
| 16.44 | 0.43 |
| 17.74 | 0.44 |
| 18.51 | 0.52 |
| 17.12 | 0.36 |
| 16.64 | 0.36 |
| 23.27 | 0.82 |
| 21.75 | 0.68 |
| 20.49 | 0.62 |
| 16.31 | 0.41 |
| 17.49 | 0.4 |

**Figure S4**. Scatter plot and linear regression of *Acanthinucella spirata* length (mm) on the x-axis and buoyant weight (g) on the y-axis (n = 629). There is a strong correlation between length and weight (p<0.001; R^2^=0.8841), validating that snail length was an accurate measure of overall snail size.

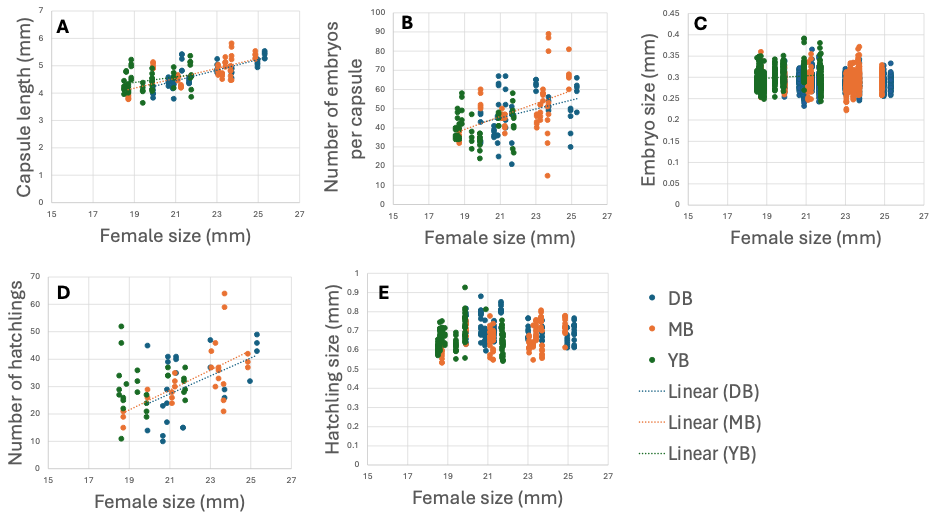

**Figure S5.** Scatter plot of average female size of *Acanthinucella spirata* on the x-axis and reproductive traits on the y-axis for **(A)** capsule length (mm), **(B)** number of embryos, **(C)** embryo diameter (mm), **(D)** number of hatchlings, and **(E)** hatchling size (mm) for Dillon Beach (DB), Muir Beach (MB) and Yerba Buena (YB). Trendlines are shown for significant (p<0.05) linear regressions (Table S12).

| Initial Length (mm) | Growth Rate (mm/day) |
| --- | --- |
| 15.9 | 0.0152597 |
| 17.8 | 0.0103896 |
| 18.2 | 0.0100649 |
| 16.2 | 0.00941558 |
| 17.7 | 0.013961 |
| 18 | 0.00227273 |
| 19.1 | 0.00779221 |
| 17.5 | 0.00357143 |
| 19.8 | 0.00324675 |
| 18 | 0.00324675 |
| 17.3 | 0.00714286 |
| 18.9 | 0.0162338 |
| 17.3 | 0.013961 |
| 18.4 | 0.00876623 |
| 16.7 | 0.00209205 |
| 17.2 | -0.00167364 |
| 16.7 | 0.00125523 |
| 17.5 | 0.0100418 |
| 17.1 | 0.0225941 |
| 17.3 | 0.0125523 |
| 18.3 | 0.0129707 |
| 16.5 | 0.00460251 |
| 18.1 | 0.00334728 |
| 17 | 0.00083682 |
| 16 | 0.00669456 |
| 18.7 | 0.00083682 |
| 18.1 | 0.0126623 |
| 19.7 | 0.012987 |
| 19.5 | 0.0117155 |
| 17.4 | 0.00962343 |
| 16.9 | 0.0142259 |
| 17 | 0.0083682 |
| 16.3 | 0.00292887 |
| 18.6 | -0.00083682 |
| 16.2 | 0.00454545 |
| 17.2 | 0.0172078 |
| 16 | 0.0185065 |
| 17.4 | 0.0136364 |
| 16 | 0.00487013 |
| 16.7 | 0.000324675 |
| 17 | 0.00909091 |
| 16.5 | 0.00487013 |
| 17.2 | 0.00487013 |
| 16.8 | 0.0136364 |
| 19.1 | 0.0142857 |
| 17.9 | 0.0194805 |
| 18 | 0.00324675 |
| 19.4 | 0.0152597 |
| 22.1 | 0.0012987 |
| 18.3 | 0.00714286 |
| 19.2 | 0.0158996 |
| 16.6 | 0.0138075 |
| 18.1 | 0.0158996 |
| 18.3 | 0.0238494 |
| 17.3 | 0.0263598 |
| 16.5 | 0.00334728 |
| 16.8 | 0.0166124 |
| 18.1 | 0.00749186 |
| 20 | 0.00879479 |
| 18.6 | 0.0224756 |
| 18.3 | 0.0019544 |
| 16.8 | 0.000651466 |
| 17.5 | 0.0149837 |
| 16.5 | 0.018241 |
| 16.4 | 0.00456026 |
| 18.4 | 0.014658 |
| 17.8 | 0.0171548 |
| 18.6 | 0.0192469 |
| 17 | 0.0224756 |
| 16.5 | 0.00130293 |
| 21.5 | 0.0112971 |
| 17.5 | 0.00543933 |
| 17.9 | 0.0251046 |
| 19.8 | 0.0200837 |
| 18.6 | 0.0337662 |
| 16.2 | 0.00519481 |
| 16.4 | 0.0165584 |
| 19.5 | 0.025 |
| 15.9 | 0.00649351 |
| 16.9 | 0.0168831 |
| 21.3 | 0.0175325 |
| 17.8 | 0.00714286 |
| 17.1 | 0.00974026 |
| 20 | 0.00422078 |
| 16 | 0.00584416 |
| 19 | 0.0191558 |
| 18.7 | 0.0230126 |
| 16.5 | 0.00669456 |
| 19.3 | 0.0175732 |
| 16.7 | -0.00125523 |
| 17.3 | 0.0129707 |
| 21.2 | 0.0117155 |
| 17.3 | 0.00714286 |
| 17 | 0.0188312 |
| 19.5 | 0.00130293 |
| 18.4 | 0.00749186 |
| 18.7 | 0.023127 |
| 16.8 | 0.00716612 |
| 18.1 | 0.013355 |
| 17.9 | 0.0166124 |
| 17.5 | 0.0107492 |
| 17.3 | 0.00228013 |
| 18.1 | 0.0211726 |
| 19.2 | 0.00684039 |
| 20.3 | 0.0166124 |
| 19.8 | 0.00456026 |
| 19.9 | 0.00488599 |
| 18.9 | 0.0208469 |
| 16.3 | 0.0041841 |
| 17 | 0.0125523 |
| 17.3 | 0.0108787 |
| 16.3 | 0.00627615 |
| 18.1 | 0.0138075 |
| 16.4 | 0.0108787 |
| 17.9 | 0.00390879 |
| 18.1 | 0.00618893 |
| 17.8 | 0 |
| 18 | 0.0167364 |
| 17.4 | 0.0133891 |
| 19.5 | 0.0083682 |
| 17.2 | 0.00460251 |
| 20 | 0.0104603 |
| 18.4 | 0.00251046 |
| 17.2 | 0.0284519 |
| 18.5 | 0.0158996 |
| 17.3 | 0.00794979 |
| 20.6 | 0.0205021 |
| 19.4 | 0.00627615 |
| 17.5 | 0.0146444 |
| 19.8 | 0.00041841 |
| 17.3 | 0.00684039 |
| 20.2 | 0.00521173 |
| 17.7 | 0.00358306 |
| 17.5 | 0.00814332 |
| 20.5 | 0.00716612 |
| 20 | 0.00684039 |
| 17.9 | 0.0276873 |
| 17.2 | 0.00553746 |
| 17.2 | 0.0175325 |
| 17.7 | 0.0133117 |
| 21.6 | 0.0107143 |
| 21.6 | 0.00714286 |
| 22.4 | 0.00811688 |
| 24.4 | 0 |
| 17.1 | 0.0178571 |
| 20.7 | 0.0243506 |
| 18.6 | 0.00941558 |
| 19.6 | 0.0107143 |
| 18.4 | 0.000649351 |
| 17 | 0.012987 |
| 19.8 | 0.0159091 |
| 16.6 | 0.00711297 |
| 18.1 | 0.00920502 |
| 18.4 | 0.0100418 |
| 16.6 | 0.0121339 |
| 17 | 0.0142259 |
| 18.5 | 0.0083682 |
| 21.8 | 0.0123377 |
| 19 | 0.00162338 |
| 17.1 | 0 |
| 17.6 | 0.00460251 |
| 19.9 | 0.0213389 |
| 16.6 | 0.00920502 |
| 19.9 | 0.00167364 |
| 20.5 | 0.0238494 |
| 24.6 | 0.00251046 |
| 18.2 | 0.00794979 |
| 21.8 | 0.00083682 |
| 21.3 | 0.00794979 |
| 22 | 0.00909091 |
| 23.5 | 0.012987 |
| 19.1 | 0.0181818 |
| 18.2 | 0.0107143 |
| 16.2 | 0.00584416 |
| 17.8 | 0.0224026 |
| 18.2 | 0.0188312 |
| 16.5 | 0.0113636 |
| 20.2 | 0.0185065 |
| 19.5 | 0.00584416 |
| 19.3 | 0.00681818 |
| 16.9 | 0.034202 |
| 16.5 | 0.0114007 |
| 19.7 | 0.0283388 |
| 20.5 | 0.0159609 |
| 18.7 | 0.0123779 |
| 19 | 0.0104235 |
| 19.2 | 0.0306189 |
| 18.4 | 0.0260586 |
| 18.5 | 0.0114007 |
| 18.7 | 0.023127 |
| 17 | 0.0221498 |
| 17.4 | 0.0247557 |
| 22.6 | 0.00456026 |
| 21 | 0.00553746 |
| 17.5 | 0.0107492 |
| 18.5 | 0.0175896 |
| 19 | 0.00920502 |
| 18.8 | 0.00585774 |
| 18.7 | 0.0117155 |
| 17.6 | 0.0322176 |

**Figure S6**. Scatter plot of initial *Acanthinucella spirata* length (mm) and average growth rate (mm/day). There was no correlation between length and growth rate (p = 0.996), so we used absolute growth rate as a response variable in our models.

**Table S1**. Table of sites used in this study, along with the latitude and region (range edge vs. core) for each site and if the site was used in the field and/or common garden experiment.

| **Site** | **Latitude** | **Region** | **Field and/or common garden experiment** |
| --- | --- | --- | --- |
| Cabrillo Beach (CB) | 33°42'31.5"N 118°17'06.1"W | Range core | Field experiment |
| Yerba Buena (YB) | 34°03'09.8"N 118°57'51.2"W | Range core | Common garden experiment |
| Rincon Point (RP) | 34°22'31.1"N 119°28'24.2"W | Range core | Field and common garden experiment |
| Muir Beach (MB) | 37°51'28.0"N 122°34'27.3"W | Range edge | Field and common garden experiment |
| Dillon Beach (DB) | 38°15'17.0"N 122°58'12.6"W | Range edge | Field and common garden experiment |
| Cape Mendocino (CM) | 40°26'09.1"N 124°24'27.9"W | Range edge | Field experiment |

**Table S2.** Metadata of *Acanthinucella spirata* breeding pairs, including initial male (M) and female (F) sizes (mm), date placed in breeding pair, and container (Cont) ID number. Population abbreviations are defined in Table S1.

| **Population** | **F_ID** | **F_Length** | **M_ID** | **M_Length** | **Cont_ID** | **Date** |  |
| --- | --- | --- | --- | --- | --- | --- | --- |
| YB | 192 | 22.1 | 468 | 18.3 | YB_H1 | 11/15/23 |  |
| YB | 201 | 19.7 | 152 | 18.1 | YB_H2 | 11/15/23 |  |
| YB | 238 | 18.4 | 166 | 17.3 | YB_H3 | 11/15/23 |  |
| YB | 419 | 18.9 | 190 | 17.3 | YB_H4 | 11/15/23 |  |
| YB | 237 | 19.8 | 434 | 18 | YB_H5 | 11/15/23 |  |
| YB | 227 | 19.1 | 247 | 17.5 | YB_H6 | 11/15/23 |  |
| YB | 481 | 18 | 259 | 17.7 | YB_H7 | 11/15/23 |  |
| YB | 210 | 18.2 | 292 | 16.2 | YB_H8 | 11/15/23 |  |
| YB | 181 | 17.8 | 141 | 15.9 | YB_H9 | 11/15/23 |  |
| YB | 489 | 19.4 | 277 | 18 | YB_H10 | 11/15/23 |  |
| YB | 447 | 17.9 | 206 | 19.1 | YB_H11 | 11/15/23 |  |
| YB | 241 | 16.8 | 140 | 17.2 | YB_H12 | 11/15/23 |  |
| YB | 122 | 17 | 255 | 16.5 | YB_H13 | 11/15/23 |  |
| YB | 261 | 16.7 | 257 | 16 | YB_H14 | 11/15/23 |  |
| YB | 429 | 17.4 | 249 | 16 | YB_H15 | 11/15/23 |  |
| YB | 252 | 17.2 | 126 | 16.2 | YB_H16 | 11/15/23 |  |
| RP | 401 | 17.3 | 441 | 17 | RP_H1 | 11/15/23 |  |
| RP | 432 | 19 | 232 | 16 | RP_H2 | 11/15/23 |  |
| RP | 458 | 20 | 245 | 17.1 | RP_H3 | 11/15/23 |  |
| RP | 254 | 21.3 | 445 | 17.8 | RP_H4 | 11/15/23 |  |
| RP | 209 | 16.9 | 81 | 15.9 | RP_H5 | 11/15/23 |  |
| RP | 95 | 19.5 | 83 | 16.4 | RP_H6 | 11/15/23 |  |
| RP | 73 | 18.6 | 272 | 16.2 | RP_H7 | 11/15/23 |  |
| MB | 124 | 19.5 | 448 | 19.3 | MB_H1 | 11/15/23 |  |
| MB | 91 | 21.8 | 115 | 19 | MB_H2 | 11/15/23 |  |
| MB | 45 | 19.8 | 282 | 18.9 | MB_H3 | 11/15/23 |  |
| MB | 156 | 17 | 4 | 18.4 | MB_H4 | 11/15/23 |  |
| MB | 101 | 18.6 | 415 | 19.6 | MB_H5 | 11/15/23 |  |
| MB | 153 | 20.7 | 44 | 17.1 | MB_H6 | 11/15/23 |  |
| MB | 37 | 22.4 | 61 | 24.4 | MB_H7 | 11/15/23 |  |
| MB | 5 | 21.6 | 197 | 21.6 | MB_H8 | 11/15/23 |  |
| MB | 168 | 17.2 | 262 | 17.7 | MB_H9 | 11/15/23 |  |
| MB | 86 | 20.2 | 133 | 17.3 | MB_H10 | 11/15/23 |  |
| MB | 479 | 18.2 | 499 | 16.5 | MB_H11 | 11/15/23 |  |
| MB | 80 | 17.8 | 43 | 16.2 | MB_H12 | 11/15/23 |  |
| MB | 77 | 19.1 | 208 | 18.2 | MB_H13 | 11/15/23 |  |
| MB | 137 | 22 | 145 | 23.5 | MB_H14 | 11/15/23 |  |
| DB | 52 | 18.2 | 82 | 19 | DB_H1 | 11/15/23 |  |
| DB | 28 | 19.5 | 193 | 18.2 | DB_H2 | 11/15/23 |  |
| DB | 50 | 23 | 246 | 20.9 | DB_H3 | 11/15/23 |  |
| DB | 296 | 19.7 | 286 | 19.4 | DB_H4 | 11/15/23 |  |
| DB | 14 | 21.1 | 271 | 20.8 | DB_H5 | 11/15/23 |  |
| DB | 273 | 19 | 15 | 17.1 | DB_H6 | 11/15/23 |  |
| DB | 105 | 20.3 | 281 | 20.2 | DB_H7 | 11/15/23 |  |
| DB | 64 | 21.6 | 49 | 22.2 | DB_H8 | 11/15/23 |  |
| DB | 76 | 17.7 | 23 | 17.7 | DB_H9 | 11/15/23 |  |
| DB | 57 | 23.7 | 110 | 22.7 | DB_H10 | 11/15/23 |  |
| DB | 98 | 19.2 | 54 | 19.4 | DB_H11 | 11/15/23 |  |
| DB | 264 | 18 | 288 | 18.7 | DB_H12 | 11/15/23 |  |
| DB | 67 | 19.3 | 480 | 20.9 | DB_H13 | 11/15/23 |  |
| DB | 119 | 18.8 | 20 | 19.5 | DB_H14 | 11/15/23 |  |
| DB | 3 | 18.8 | 26 | 19.5 | DB_H15 | 11/15/23 |  |
| DB | 11 | 17.8 | 46 | 17.6 | DB_H16 | 11/15/23 |  |
| DB | 275 | 19.2 | 29 | 17.8 | DB_H17 | 11/15/23 |  |
| DB | 459 | 18.3 | 39 | 17.3 | DB_H18 | 11/15/23 |  |
| DB | 196 | 19 | 199 | 20.1 | DB_H19 | 11/15/23 |  |
| YB | 527 | 17 | 639 | 16.5 | YB_C1 | 11/16/23 |  |
| YB | 646 | 18.4 | 437 | 16.4 | YB_C2 | 11/16/23 |  |
| YB | 283 | 17.5 | 677 | 16.5 | YB_C3 | 11/16/23 |  |
| YB | 300 | 18.3 | 462 | 16.8 | YB_C4 | 11/16/23 |  |
| YB | 689 | 18.6 | 478 | 20 | YB_C5 | 11/16/23 |  |
| YB | 616 | 18.1 | 405 | 16.8 | YB_C6 | 11/16/23 |  |
| RP | 636 | 17.9 | 652 | 17.2 | RP_C1 | 11/16/23 |  |
| RP | 559 | 18.1 | 336 | 17.9 | RP_C2 | 11/16/23 |  |
| RP | 506 | 18.9 | 502 | 19.9 | RP_C3 | 11/16/23 |  |
| RP | 627 | 20.3 | 687 | 19.8 | RP_C4 | 11/16/23 |  |
| RP | 577 | 18.1 | 580 | 19.2 | RP_C5 | 11/16/23 |  |
| RP | 678 | 17.3 | 510 | 17.5 | RP_C6 | 11/16/23 |  |
| RP | 600 | 18.1 | 694 | 17.9 | RP_C7 | 11/16/23 |  |
| RP | 693 | 16.8 | 575 | 18.7 | RP_C8 | 11/16/23 |  |
| RP | 683 | 18.4 | 560 | 19.5 | RP_C9 | 11/16/23 |  |
| RP | 514 | 20.5 | 666 | 20 | RP_C10 | 11/16/23 |  |
| RP | 393 | 17.7 | 650 | 17.5 | RP_C11 | 11/16/23 |  |
| RP | 380 | 17.3 | 507 | 20.2 | RP_C12 | 11/16/23 |  |
| MB | 596 | 19 | 303 | 20.2 | MB_C1 | 11/16/23 |  |
| MB | 523 | 18.5 | 304 | 17.5 | MB_C2 | 11/16/23 |  |
| MB | 581 | 21 | 533 | 22.6 | MB_C3 | 11/16/23 |  |
| MB | 592 | 17.4 | 525 | 17 | MB_C4 | 11/16/23 |  |
| MB | 550 | 18.7 | 537 | 18.5 | MB_C5 | 11/16/23 |  |
| MB | 394 | 19.2 | 551 | 18.4 | MB_C6 | 11/16/23 |  |
| MB | 505 | 18.7 | 530 | 19 | MB_C7 | 11/16/23 |  |
| MB | 305 | 19.7 | 362 | 20.5 | MB_C8 | 11/16/23 |  |
| MB | 503 | 16.9 | 532 | 16.5 | MB_C9 | 11/16/23 |  |
| DB | 509 | 17.4 | 520 | 17.3 | DB_C1 | 11/16/23 |  |
| DB | 554 | 18.9 | 534 | 18.9 | DB_C2 | 11/16/23 |  |
| DB | 688 | 19 | 595 | 20 | DB_C3 | 11/16/23 |  |
| DB | 571 | 18.2 | 425 | 19 | DB_C4 | 11/16/23 |  |
| DB | 495 | 17.6 | 667 | 19.1 | DB_C5 | 11/16/23 |  |
| DB | 526 | 18.8 | 591 | 17.7 | DB_C6 | 11/16/23 |  |
| DB | 512 | 18.3 | 607 | 17.2 | DB_C7 | 11/16/23 |  |
| DB | 570 | 17.8 | 565 | 16.5 | DB_C8 | 11/16/23 |  |
| DB | 327 | 18.3 | 542 | 16.5 | DB_C9 | 11/16/23 |  |
| DB | 508 | 17.1 | 521 | 17.9 | DB_C10 | 11/16/23 |  |
| DB | 638 | 17.9 | 547 | 18.2 | DB_C11 | 11/16/23 |  |
| DB | 426 | 17.9 | 411 | 16.7 | DB_C12 | 11/16/23 |  |
| DB | 540 | 17.7 | 597 | 18 | DB_C13 | 11/16/23 |  |
| YB | 568 | 17.3 | 609 | 16.5 | YB_C7 | 1/23/24 |  |
| YB | 544 | 18.3 | 170 | 18.1 | YB_C8 | 1/23/24 |  |
| YB | 114 | 19.2 | 511 | 16.6 | YB_C9 | 1/23/24 |  |
| YB | 490 | 18.6 | 471 | 17.8 | YB_C10 | 1/23/24 |  |
| RP | 585 | 19.8 | 309 | 17.5 | RP_C13 | 1/23/24 |  |
| RP | 333 | 20.6 | 663 | 19.4 | RP_C14 | 1/23/24 |  |
| RP | 583 | 18.5 | 647 | 17.3 | RP_C15 | 1/23/24 |  |
| RP | 685 | 17.2 | 588 | 18.4 | RP_C16 | 1/23/24 |  |
| RP | 366 | 17.2 | 513 | 20 | RP_C17 | 1/23/24 |  |
| RP | 331 | 17.4 | 699 | 19.5 | RP_C18 | 1/23/24 |  |
| RP | 621 | 18 | 538 | 17.8 | RP_C19 | 1/23/24 |  |
| RP | 624 | 16.4 | 322 | 18.1 | RP_C20 | 1/23/24 |  |
| RP | 350 | 16.3 | 342 | 17.3 | RP_C21 | 1/23/24 |  |
| RP | 373 | 16.3 | 605 | 17 | RP_C22 | 1/23/24 |  |
| MB | 528 | 25 | 387 | 19.5 | MB_C10 | 1/23/24 |  |
| MB | 396 | 17.6 | 361 | 18.7 | MB_C11 | 1/23/24 |  |
| MB | 582 | 19 | 590 | 18.8 | MB_C12 | 1/23/24 |  |
| DB | 553 | 19.1 | 567 | 18.3 | DB_C14 | 1/23/24 |  |
| DB | 374 | 20.1 | 320 | 16.9 | DB_C15 | 1/23/24 |  |
| DB | 656 | 19 | 641 | 16.3 | DB_C16 | 1/23/24 |  |
| YB | 219 | 18.6 | 125 | 16.3 | YB_H17 | 1/23/24 |  |
| YB | 235 | 16.9 | 251 | 17 | YB_H18 | 1/23/24 |  |
| YB | 175 | 19.5 | 293 | 17.4 | YB_H19 | 1/23/24 |  |
| YB | 270 | 18.7 | 180 | 16 | YB_H20 | 1/23/24 |  |
| YB | 177 | 18.1 | 213 | 17 | YB_H21 | 1/23/24 |  |
| YB | 221 | 18.3 | 474 | 16.5 | YB_H22 | 1/23/24 |  |
| YB | 147 | 17.3 | 113 | 17.1 | YB_H23 | 1/23/24 |  |
| YB | 443 | 17.5 | 233 | 16.7 | YB_H24 | 1/23/24 |  |
| YB | 260 | 17.2 | 222 | 16.7 | YB_H25 | 1/23/24 |  |
| RP | 453 | 19.8 | 276 | 17.9 | RP_H8 | 1/23/24 |  |
| RP | 299 | 21.5 | 414 | 17.5 | RP_H9 | 1/23/24 |  |
| RP | 266 | 21.2 | 212 | 17.3 | RP_H10 | 1/23/24 |  |
| RP | 72 | 19.3 | 436 | 16.7 | RP_H11 | 1/23/24 |  |
| RP | 229 | 18.7 | 465 | 16.5 | RP_H12 | 1/23/24 |  |
| MB | 108 | 21.8 | 135 | 21.3 | MB_H15 | 1/23/24 |  |
| MB | 96 | 24.6 | 188 | 18.2 | MB_H16 | 1/23/24 |  |
| MB | 128 | 20.5 | 2 | 19.9 | MB_H17 | 1/23/24 |  |
| MB | 100 | 19.9 | 134 | 16.6 | MB_H18 | 1/23/24 |  |
| MB | 042* | 17.1 | 169 | 17.6 | MB_H19 | 1/23/24 | *042 replaced 060 on 2/27/24 |
| MB | 223 | 18.5 | 187 | 17 | MB_H20 | 1/23/24 |  |
| MB | 70 | 18.4 | 123 | 16.6 | MB_H21 | 1/23/24 |  |
| MB | 144 | 18.1 | 75 | 16.6 | MB_H22 | 1/23/24 |  |
| DB | 171 | 18.9 | 172 | 17.6 | DB_H20 | 1/23/24 |  |
| DB | 189 | 17.5 | 35 | 16.3 | DB_H21 | 1/23/24 |  |
| DB | 22 | 17.2 | 12 | 17 | DB_H22 | 1/23/24 |  |
| DB | 211 | 30.2 | 269 | 18.6 | DB_H23 | 1/23/24 |  |

**Table S3.** Pairwise comparisons from contrast function in the *emmeans* package for capsule length from field-collected *A. spirata* egg capsules. Significant pairwise comparisons (p<0.05) are in bold. Population abbreviations are defined in Table S1.

| **contrast** | **estimate** | **SE** | **df** | **t.ratio** | **p-value** |
| --- | --- | --- | --- | --- | --- |
| CB - CM | -0.2914 | 0.0315 | 87 | -9.239 | **<0.0001** |
| CB - DB | -0.24 | 0.0306 | 87 | -7.838 | **<0.0001** |
| CB - MB | -0.1081 | 0.0315 | 87 | -3.429 | **0.0081** |
| CB - RP | -0.1239 | 0.0391 | 87 | -3.17 | **0.0175** |
| CM - DB | 0.0513 | 0.0261 | 87 | 1.966 | 0.2912 |
| CM - MB | 0.1832 | 0.0272 | 87 | 6.741 | **<0.0001** |
| CM - RP | 0.1674 | 0.0357 | 87 | 4.694 | **0.0001** |
| DB - MB | 0.1319 | 0.0261 | 87 | 5.05 | **<0.0001** |
| DB - RP | 0.1161 | 0.0349 | 87 | 3.329 | **0.0109** |
| MB - RP | -0.0158 | 0.0357 | 87 | -0.443 | 0.9919 |

**Table S4.** Pairwise comparisons from contrast function in the *emmeans* package for number of embryos per capsule from field-collected *A. spirata* egg capsules. Significant pairwise comparisons (p<0.05) are in bold. Population abbreviations are defined in Table S1.

| **contrast** | **estimate** | **SE** | **df** | **z.ratio** | **p.value** |
| --- | --- | --- | --- | --- | --- |
| CB - CM | -0.4046 | 0.0796 | Inf | -5.081 | **<0.0001** |
| CB - DB | -0.481 | 0.0772 | Inf | -6.227 | **<0.0001** |
| CB - MB | -0.3829 | 0.0797 | Inf | -4.802 | **<0.0001** |
| CB - RP | -0.0633 | 0.1011 | Inf | -0.626 | 0.9709 |
| CM - DB | -0.0764 | 0.0614 | Inf | -1.243 | 0.7258 |
| CM - MB | 0.0216 | 0.0646 | Inf | 0.335 | 0.9973 |
| CM - RP | 0.3413 | 0.0896 | Inf | 3.807 | **0.0013** |
| DB - MB | 0.098 | 0.0616 | Inf | 1.591 | 0.5028 |
| DB - RP | 0.4177 | 0.0875 | Inf | 4.772 | **<0.0001** |
| MB - RP | 0.3196 | 0.0897 | Inf | 3.561 | **0.0034** |

**Table S5.** Pairwise comparisons from contrast function in the *emmeans* package for embryo size from field-collected *A. spirata* egg capsules. Significant pairwise comparisons (p<0.05) are in bold. Population abbreviations are defined in Table S1.

| **contrast** | **estimate** | **SE** | **df** | **t.ratio** | **p.value** |
| --- | --- | --- | --- | --- | --- |
| CB - CM | 0.01282 | 0.007 | 22.9 | 1.833 | 0.3800 |
| CB - DB | 0.01953 | 0.00678 | 21.6 | 2.881 | 0.0604 |
| CB - MB | 0.02153 | 0.00699 | 23 | 3.083 | **0.0381** |
| CB - RP | 0.02943 | 0.00861 | 26.7 | 3.419 | **0.0159** |
| CM - DB | 0.00671 | 0.00596 | 18.3 | 1.127 | 0.7906 |
| CM - MB | 0.00871 | 0.00619 | 20 | 1.408 | 0.6300 |
| CM - RP | 0.0166 | 0.00797 | 25 | 2.082 | 0.2587 |
| DB - MB | 0.002 | 0.00594 | 18.5 | 0.336 | 0.9970 |
| DB - RP | 0.00989 | 0.00778 | 24.1 | 1.271 | 0.7109 |
| MB - RP | 0.00789 | 0.00796 | 25.2 | 0.991 | 0.8568 |

**Table S6.** Pairwise comparisons from contrast function in the *emmeans* package for nurse egg ratio from field-collected *A. spirata* egg capsules. Significant pairwise comparisons (p<0.05) are in bold. Population abbreviations are defined in Table S1.

| **contrast** | **estimate** | **SE** | **df** | **t.ratio** | **p.value** |
| --- | --- | --- | --- | --- | --- |
| CB - CM | -15.39 | 6.35 | 17 | -2.426 | 0.1559 |
| CB - DB | -22.48 | 4.44 | 17 | -5.061 | **0.0008** |
| CB - MB | -20.76 | 4.8 | 17 | -4.327 | **0.0036** |
| CB - RP | -5.97 | 5.54 | 17 | -1.077 | 0.8157 |
| CM - DB | -7.09 | 6.08 | 17 | -1.165 | 0.7703 |
| CM - MB | -5.37 | 6.35 | 17 | -0.846 | 0.9124 |
| CM - RP | 9.43 | 6.92 | 17 | 1.362 | 0.6587 |
| DB - MB | 1.72 | 4.44 | 17 | 0.387 | 0.9948 |
| DB - RP | 16.51 | 5.23 | 17 | 3.155 | **0.0401** |
| MB - RP | 14.79 | 5.54 | 17 | 2.671 | 0.1010 |

**Table S7.** Pairwise comparisons from contrast function in the *emmeans* package for hatchling size from field-collected *A. spirata* egg capsules. Significant pairwise comparisons (p<0.05) are in bold. Population abbreviations are defined in Table S1.

| **contrast** | **estimate** | **SE** | **df** | **t.ratio** | **p.value** |
| --- | --- | --- | --- | --- | --- |
| CB - CM | -0.070686 | 0.012 | 467 | -5.873 | **<0.0001** |
| CB - DB | -0.08312 | 0.0126 | 467 | -6.581 | **<0.0001** |
| CB - MB | -0.015137 | 0.0136 | 467 | -1.111 | 0.8007 |
| CB - RP | -0.015641 | 0.0123 | 467 | -1.274 | 0.7075 |
| CM - DB | -0.012435 | 0.0115 | 467 | -1.086 | 0.8139 |
| CM - MB | 0.055548 | 0.0125 | 467 | 4.43 | **0.0001** |
| CM - RP | 0.055045 | 0.0111 | 467 | 4.976 | **<0.0001** |
| DB - MB | 0.067983 | 0.0131 | 467 | 5.185 | **<0.0001** |
| DB - RP | 0.06748 | 0.0117 | 467 | 5.763 | **<0.0001** |
| MB - RP | -0.000503 | 0.0128 | 467 | -0.039 | 1.000 |

**Table S8.** Pairwise comparisons from the contrast function in the *emmeans* package for embryo size from the A. *spirata* egg capsules laid in the common garden experiment. Significant pairwise comparisons (p<0.05) are in bold. Population abbreviations are defined in Table S1.

| **contrast** | **estimate** | **SE** | **df** | **t.ratio** | **p.value** |
| --- | --- | --- | --- | --- | --- |
| DB - MB | -0.0022 | 0.0038 | 26 | -0.578 | 0.8328 |
| DB - YB | -0.01073 | 0.00416 | 26 | -2.577 | **0.0410** |
| MB - YB | -0.00853 | 0.00418 | 26 | -2.038 | 0.1231 |

**Table S9.** Pairwise comparisons from the contrast function in the *emmeans* package for hatchling size from the A. *spirata* egg capsules laid in the common garden experiment. Significant pairwise comparisons (p<0.05) are in bold. Population abbreviations are defined in Table S1.

| **contrast** | **etimate** | **SE** | **df** | **t.ratio** | **p.value** |
| --- | --- | --- | --- | --- | --- |
| DB - MB | 0.05046 | 0.012 | 394 | 4.195 | **0.0001** |
| DB - YB | 0.04638 | 0.0135 | 394 | 3.424 | **0.0020** |
| MB - YB | -0.00408 | 0.0135 | 394 | -0.303 | 0.9507 |

**Table S10.** Pairwise comparisons from the contrast function in the *emmeans* package for capsule length from the A. *spirata* egg capsules laid in the common garden experiment. Significant pairwise comparisons (p<0.05) are in bold. Population abbreviations are defined in Table S1.

| **contrast** | **estimate** | **SE** | **df** | **t.ratio** | **p.value** |
| --- | --- | --- | --- | --- | --- |
| DB - MB | -0.0163 | 0.0152 | 146 | -1.073 | 0.5322 |
| DB - YB | -0.0439 | 0.0178 | 146 | -2.472 | **0.0386** |
| MB - YB | -0.0275 | 0.0179 | 146 | -1.54 | 0.2752 |

**Table S11.** Pairwise comparisons from the contrast function in the *emmeans* package for fecundity from the A. *spirata* egg capsules laid in the common garden experiment. Significant pairwise comparisons (p<0.05) are in bold. Population abbreviations are defined in Table S1.

| **contrast** | **estimate** | **SE** | **df** | **t.ratio** | **p.value** |
| --- | --- | --- | --- | --- | --- |
| DB - MB | -0.0818 | 0.213 | 25 | -0.384 | 0.9223 |
| DB - YB | -0.5224 | 0.209 | 25 | -2.499 | **0.0491** |
| MB - YB | -0.4406 | 0.212 | 25 | -2.079 | 0.1147 |

**Table S12.** Linear regression significance values for the correlation between snail length (mm) and reproductive traits for each population in the common garden experiment. Significant correlations (p<0.05) are in bold. Population abbreviations are defined in Table S1.

| Site | Capsule Length | Number of embryos | Embryo Size | Number of hatchlings | Hatchling size |
| --- | --- | --- | --- | --- | --- |
| YB | **0.027** | 0.332 | <0.001 | 0.776 | 0.923 |
| MB | **<0.001** | **0.0012** | 0.642 | **<0.001** | <0.001 |
| DB | **<0.001** | **0.020** | 0.239 | **0.006** | 0.467 |
